# Exploring the Persistence of Transgenes in Genetically Engineered Cyanobacteria

**DOI:** 10.1101/2025.11.26.690176

**Authors:** Cherrelle L. Barnes, James W. Lee, Lesley H. Greene

## Abstract

Genetically engineered organisms including bacteria, plants, and animals are very commonplace in industry and scientific research where they are designed to perform specific tasks. Cyanobacteria are a group of ubiquitous and ancient microorganisms that have been engineered to produce a range of products such as biofuels. However, the potential ramifications of genetic engineering could have unintended consequences. One key question is how long do foreign genes persist? Thus, we undertook a two-year study that investigates the fate and stability of transgenes in genetically engineered *Thermosynechococcus elongatus* BP1. The results show transgenes are very persistent within the host genome and begin to become lost slowly over time.

## Introduction

Transgene stability can be defined as the persistent presence or expression of a foreign gene within a host. For commercial and scientific use of genetically engineered (GE) organisms, the goal is for the organisms to possess and express their transgenes for the duration of their lifetime. Transgene stability has been studied in many GE organisms including bacteria, plants, and insects (Handler, 2004; Li et al., 2009; Abidin et al., 2021). Transgene stability within GE organisms can be affected through gene silencing by epigenetic and transcriptional mechanisms or gene loss because of crossbreeding (Finnegan and McElroy, 1994; Dietz-Pfeilstetter, 2010). From a bio-safety perspective, a concern about the development of GE organisms is the fate and stability of transgenes within the host, as it could lead to gene escape and allow transgenes to persist in unintended environments (Warwick et al., 2008; Agga et al., 2019). It is important to investigate and understand long-term stability of transgenes, as it could have an impact on the environment as well as human and animal health. The research presented in this study focuses on assessing the fate and stability of transgenes within genetically engineered cyanobacteria, which can be of great concern if they escape containment. It is also of importance to know how long transgenes persist in GE organisms to address general mechanisms in microbial genetics and evolution.

To gain a better understanding of the outcome and long-term stability of transgenes inserted into host chromosomes, we conducted an exploratory study (Barnes, 2022). Here cultures of a model cyanobacteria with a cassette of transgenes which included the kanamycin resistance gene as an antibiotic selection marker were incubated with and without antibiotic for a two-year study. Monthly, genomic material was extracted from each culture and used to determine the presence of the transgene cassette which revealed significant stability as evidenced from their persistence in the genome.

## Results and Discussion

### The presence of the pUC57-pKA cassette slowly decreases over time without antibiotic pressure

Each month genomic DNA was isolated and PCR of the insertion site, the *Kan* resistance gene and the *rpsL* gene, which served as a control, was conducted to determine the presence of the integrated transgenes over a two-year period. PCR was used to amplify the insert region (Figure 1) for the transgenes, resulting in a 6.7 kb or 2.4 kb band indicating chromosome copies that have or have not integrated the transgene cassette, respectively. Initially, the 2.4 kb bands are much less intense than the 6.7 kb bands among all cultures, with and without antibiotic selective pressure (Figure 2). This indicated that a higher number of chromosomes within the culture have integrated transgenes. This trend remained the same for cultures with antibiotic selective pressure for the entire duration of the study. However, in the cultures without antibiotic selective pressure, the 2.4 kb band become more intense indicating that a portion of the chromosomes with integrated transgenes decreases over time (Figure 2). This trend is evident in the two-year stability study.

**Figure 1.**
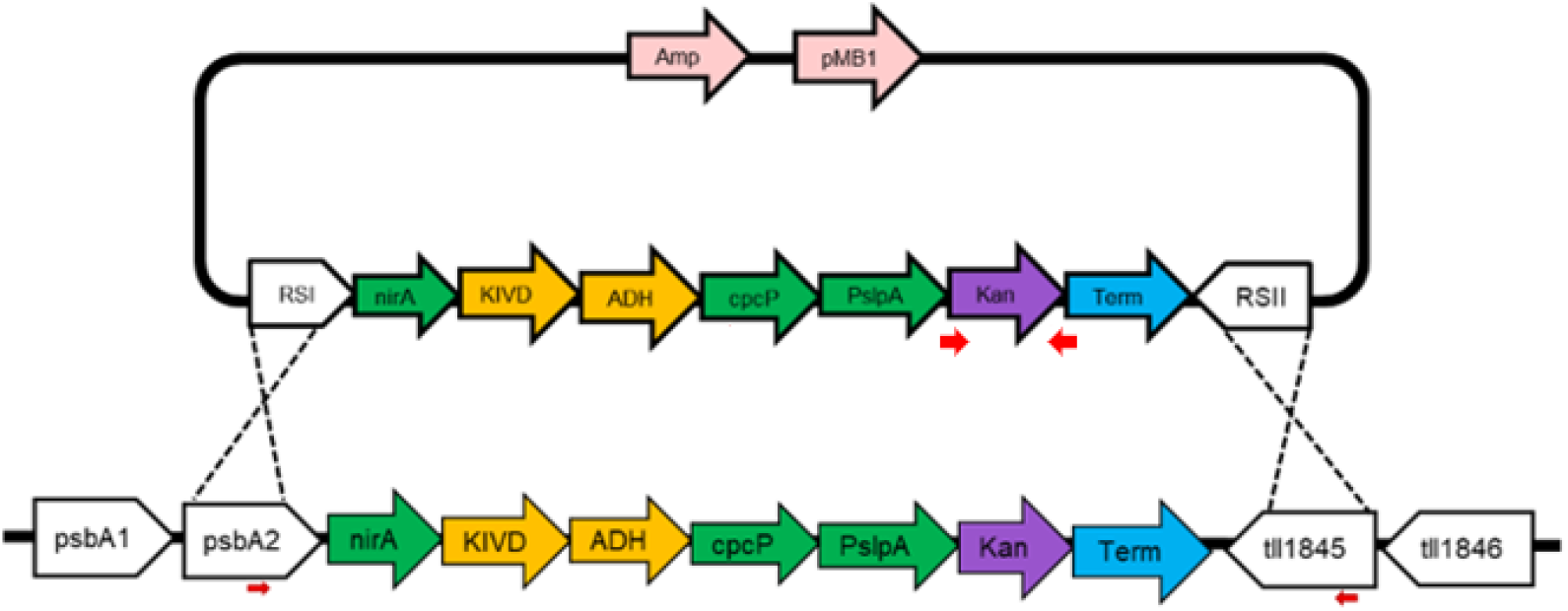
Schematic of insertion and PCR sites. Primers depicted as red arrows were used to amplify the kanamycin resistance gene and the insertion site within the genome. KIVD = alpha-ketoisovalerate decarboxylase, ADH = NADH-dependent alcohol dehydrogenase, Kan = Kanamycin resistance gene.

**Figure 2.**
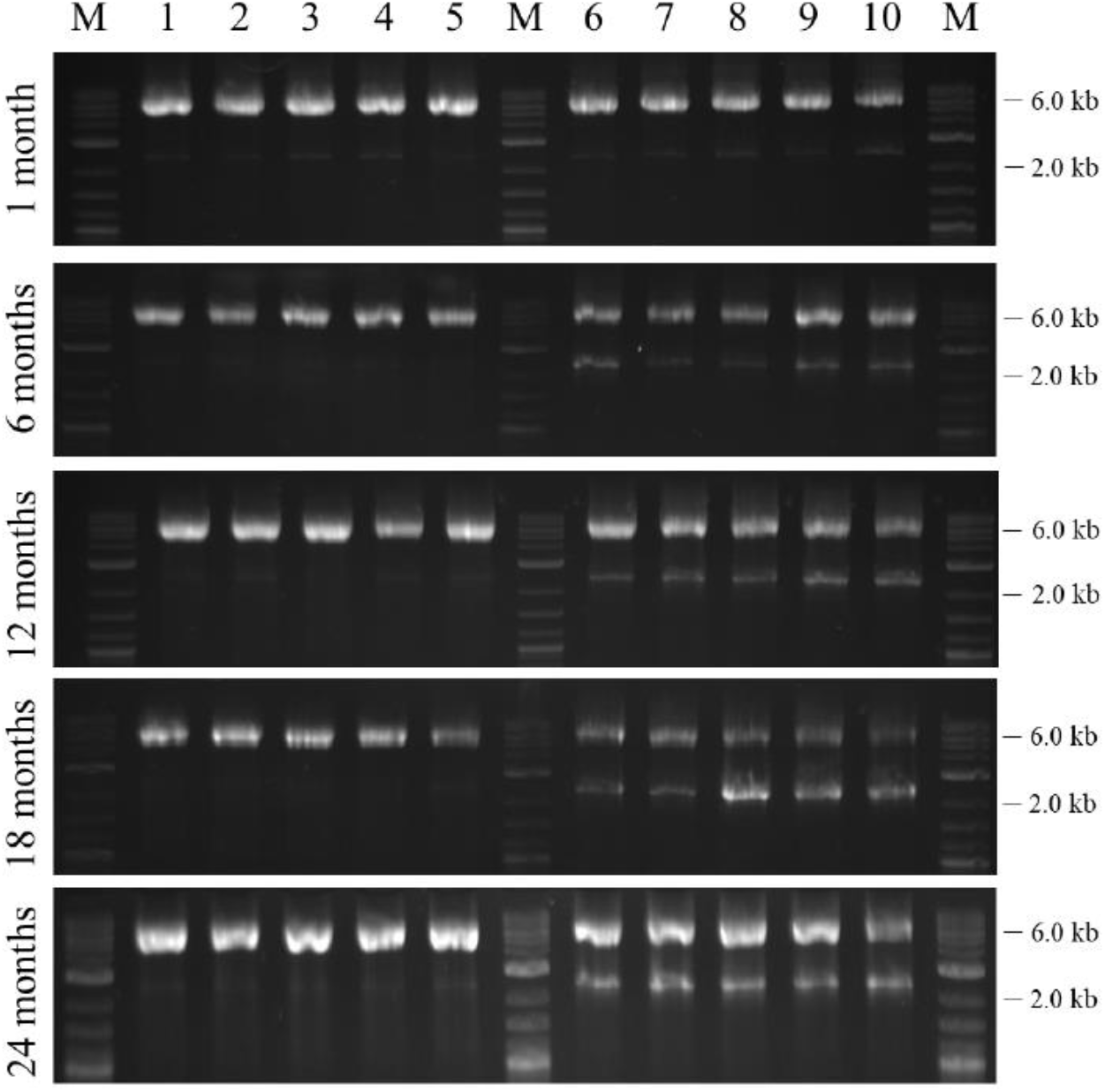
Presence of the gene cassette within GE *T. elongatus* BP1 over 24 months. Primers were used to amplify the insertion site of *T. elongatus* BP1 genome containing the gene cassette, resulting in either a 2.4 kb band (wild-type size, no integration of genes) or 6.7 kb band (integration of genes) at 1, 6, 12, 18 and 24 months. The lanes are marked as followed: M = molecular weight marker, 1-5 = genomic DNA from the control GE *T. elongatus* BP1-pKA cultures (+ Kan) and 6-10 = genomic DNA from the experimental GE *T. elongatus* BP1-pKA cultures (- Kan).

To further assess these results, there was a control PCR reaction which amplified the *Kan* resistance gene resulting in a 0.7 kb band. The *Kan* resistance gene remained present within the genomic DNA of GE *T. elongatus* BP1-pKA for the two-year period (Figure 3). Further, the kanamycin resistance gene was shown to still be expressed after one year of study, even in the absence of antibiotic pressure (data not shown). One important aspect of stable transgene expression within GE organisms is the choice for the promoter. The transgene cassette was designed where the expression of the kanamycin resistance gene was under the control of the *cpc* and *slpA* continuous promoters, which are known to be very strong (Hynönen et al., 2010; Zhou et al., 2014).

**Figure 3.**
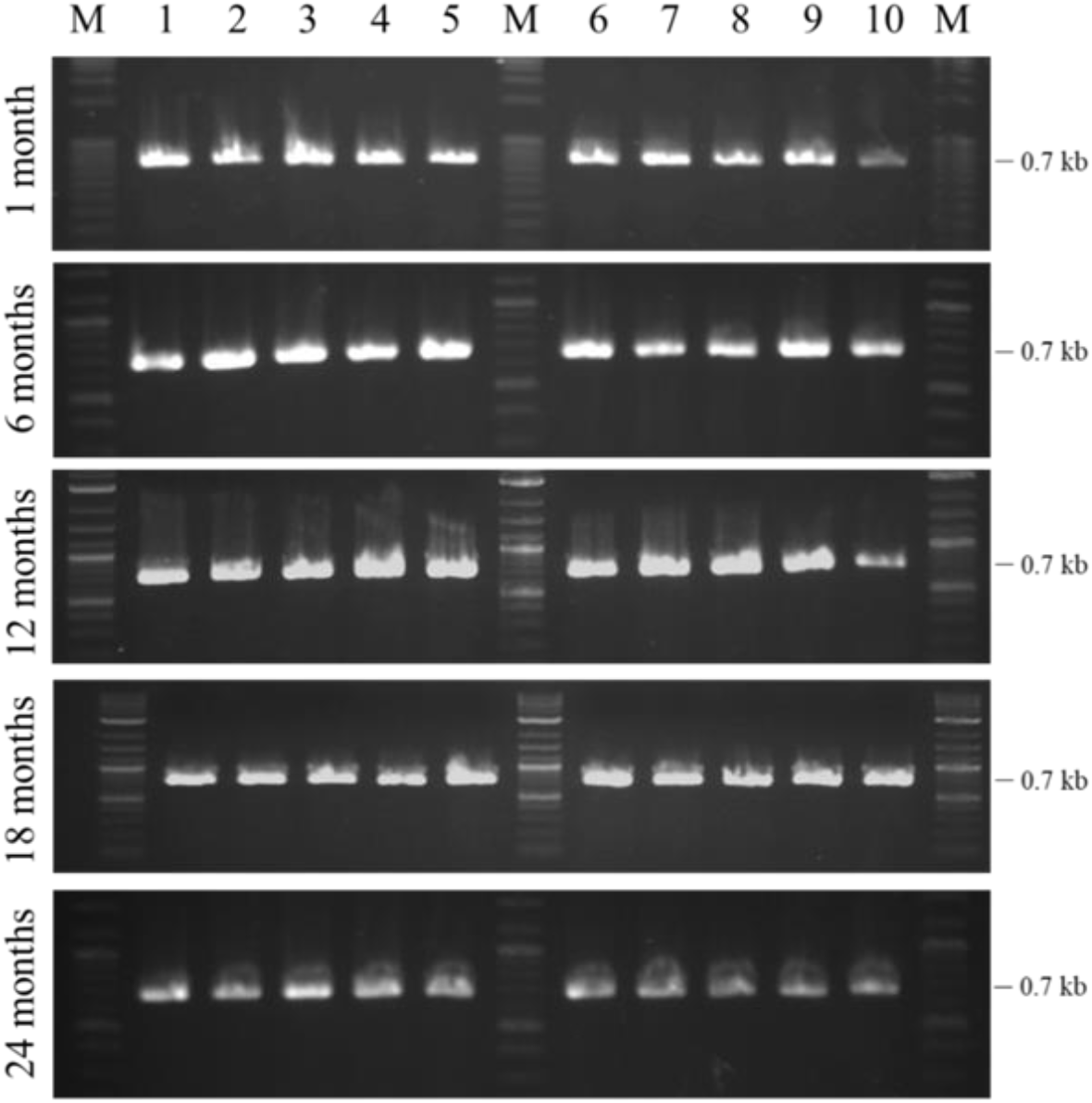
Control for the presence of the *Kan* resistance gene within GE *T. elongatus* BP1 for 24 months. Primers were used to amplify the *Kan* resistance gene within the gene cassette, resulting in a 0.7 kb band at 1, 6, 12, 18 and 24 months. The lanes are marked as followed: M = molecular weight marker, 1-5 = genomic DNA from the control GE *T. elongatus* BP1-pKA cultures (+ Kan) and 6-10 = genomic DNA from the experimental GE *T. elongatus* BP1-pKA cultures (- Kan).

## Conclusions

A gradual decrease in the population of chromosomes that have the transgene cassette was observed over the two-year study with genomic DNA. One explanation for the loss of genes over time could be genome streamlining. Bacteria, including cyanobacteria, have been known to reduce the size of the chromosomes as an evolutionary mechanism, resulting in the loss of genes not essential for survival of the organism (Giovannoni et al., 2005; Marais et al., 2008; Moya et al., 2009; Yus et al., 2009). Genomic streamlining has been demonstrated within *Prochlorococcus marinus* and *Pelagibacter ubique*, which both have genome sizes around or less than 2 Mb but still possesses essential genes for biosynthesis of all amino acids and complete metabolic network (Giovannoni et al., 2005; Moya et al., 2009). Genomic studies have also revealed evidence that several strains of the cyanobacterium *Phlanktothrix* lost the microcystin synthetase gene cluster through evolutionary deletion events, resulting in the strains becoming nontoxic (Christiansen et al., 2008). Although the results in the genomic DNA PCR show there is loss of the transgene cassette within the GE *T. elongatus* BP1 over time the specific mechanism of this gene loss is uncertain.

In general, the genomic stability of the transgene cassette is shown to be very stable for up to two years within the chromosome of *T. elongatus* BP1, both in the presence and absence of antibiotic pressure. This finding was very surprising as we initially theorized the transgenes would be lost more rapidly without the selective pressure of kanamycin in the media. There have been a few previous studies that investigated long-term transgene stability within plants and fungus (Weaver et al., 2005; Li et al., 2009; Zeng et al., 2009). One group investigated the long-term stability and expression of the *rolC* gene within GE aspen trees, which revealed that the transgene was still present within the genome and expression up to 18 years after transformation (Li et al., 2009). A different group examined the stability of transgenes within GE *Trichoderma virens* over a 250-day experiment and found that the *opd* gene was still present and express in the absence of antibiotic pressure (Weaver et al., 2005). Another study involved assessing the expression of a chimeric gene, including an insecticidal peptide gene and the C-peptide of *Bt* gene, within *Betula platyphylla* for up to 15 subcultures and although they found the gene was silenced, it was still present within the chromosome for the duration of the study (Zeng et al., 2009). There have been no long-term studies of transgene stability in cyanobacteria previously published based on extensive literature searches.

From a bio-risk perspective, the longer transgenes remain intact within the cyanobacterial chromosome, the more opportunity it may have to transfer nonnative genes and persist in unintended environments, thus leading to possible ecological and human/animal health risks. For example, a study assessed the persistence of an escaped herbicide resistance gene from GE *Brassica rapa* to wild-type relatives through hybridization and the transgene was shown to be persistent in these hybrids for up to 6 years (Warwick et al., 2008). Another study demonstrated that antibiotic resistance genes of bacteria within beef cattle is shown to be transferred and persists for up to two years in feeding areas where they were housed (Agga et al., 2019). If transgenic cyanobacteria are to be used for the commercial production of bioproducts, it is of paramount importance to understand the fate and stability of transgenes as an avenue to prevent any bio-risk concerns. Thus, it is critical that science looks ahead and seeks to understand the outcome of foreign genes.

## Materials and Methods

### Growth studies

A GE *Thermosynechococcus elongatus* BP1 (called GE T. elongatus BP-pKA) carrying a cassette of synthetic transgenes including the kanamycin resistance gene in its chromosome (Nguyen et al., 2019) was incubated in the presence and absence of antibiotic selection pressure. PCR was performed to monitor the inserted transgenes for up to 104 weeks (2 years). GE *T. elongatus* BP1 cells were grown in BG-11 liquid media with a pH ∼7.75. One triplicate set of cultures was supplemented with kanamycin 40 µg/mL to represent the presence of selective pressure and will act as the control for this study, the other triplicate set did not contain any antibiotic to represent absence of selective pressure and will act as the experimental cultures. Growth occurred in a Percival environmental chamber at ∼42 °C with continuous illumination at actinic light intensity of 30 µE m^-2^ s^-1^. The cultures were re-inoculated bi-weekly into fresh BG-11 media (3 mL of culture into 75 mL of media), and 2 mL aliquots of each culture were collected and cryopreserved monthly. The pH of the cultures was measured average at the start of incubation and following 2 weeks of growth prior to reinoculation into fresh media and the pH was ∼7.91 and ∼9.49, respectively.

### Detection of transgene cassette by PCR

Genomic DNA was extracted monthly using the Qiagen QiaAMP DNA Mini Kit and used to amplify two genetic regions: the insertion site and the *Kan* resistance gene via PCR using 2 ng of purified genomic DNA. For the insertion site, primers were designed to amplify from within the upstream recombination site of designer transgene cassette (homologous to *T. elongatus* BP1) to outside the downstream recombination site on the chromosome (not part of transgene cassette) (Figure 1). *T. elongatus* BP1 possess multiple copies of their chromosome, resulting at the onset of the study in some copies that contain the transgenes while possibly others do not. Because of this, there are two expected band sizes for the PCR amplified insert and are located at 2.4 kb and 6.7 kb on an agarose gel, which represents no integration and integration of the transgenes, respectively. The *Kan* resistance gene was also amplified and were detectable as 0.7 kb. The presence of genomic DNA was tested monthly over a two-year period to monitor any changes over time.

## Acknowledgements

This work is supported by Biotechnology Risk Assessment Grant Program competitive grant award no. 2016-33522-25624 and 2023-33522-40974 from the U.S. Department of Agriculture to JWL and LHG and support from Old Dominion University to LHG.

## References

Abidin AAZ, Othman NA, Yusoff FM, Yusof, ZNB. 2021. Determination of transgene stability in Nannochloropsis sp. transformed with immunogenic peptide for oral vaccination against vibriosis. Aquac Int 29: 477–486.

Agga GE, Cook KL, Netthisinghe AMP, Gilfillen RA, Woosley PB, Sistani KR. 2019. Persistence of antibiotic resistance genes in beef cattle backgrounding environment over two years after cessation of operation. PLoS One 14: e0212510.

Barnes, CL. 2022. Investigating the Biorisk of Genetically Engineered Thermosynechococcus Elongatus BP1. Dissertation, Old Dominion University, Norfolk, VA USA.

Christiansen G, Molitor C, Philmus B, Kurmayer R. 2008. Nontoxic Strains of Cyanobacteria Are the Result of Major Gene Deletion Events Induced by a Transposable Element. Mol Biol Evol 25: 1695–1704.

Dietz-Pfeilstetter A. 2010. Stability of transgene expression as a challenge for genetic engineering. Plant Sci 179: 164–167.

Finnegan J, McElroy D. 1994. Transgene Inactivation: Plants Fight Back! Nat Biotechnol 12: 883–888.

Giovannoni SJ, Tripp HJ, Givan S, Podar M, Vergin KL, Baptista D, Bibbs L, Eads J, Richardson TH, Noordewier M, et al. 2005. Genome streamlining in a cosmopolitan oceanic bacterium. Science 309: 1242–1245.

Handler AM. 2004. Understanding and improving transgene stability and expression in insects for SIT and conditional lethal release programs. Insect Biochem Mol Biol 34: 121–130.

Hynönen U, Avall-Jääskeläinen S, Palva, A. 2010. Characterization and separate activities of the two promoters of the Lactobacillus brevis S-layer protein gene. Appl Microbiol Biotechnol 87: 657–668.

Li J, Brunner AM, Meilan R, Strauss SH. 2009. Stability of transgenes in trees: expression of two reporter genes in poplar over three field seasons. Tree Physiol 29: 299–312.

Marais GA, Calteau A, Tenaillon O. 2008. Mutation rate and genome reduction in endosymbiotic and free-living bacteria. Genetica 134: 205–210.

Moya A, Gil R, Latorre A, Peretó J, Pilar Garcillán-Barcia M, De La Cruz, F. 2009. Toward minimal bacterial cells: evolution vs. design. FEMS Microbiol Rev 33: 225–235.

Nguyen TH, Barnes CL, Agola JP, Sherazi S, Greene LH, Lee JW. 2019. Demonstration of horizontal gene transfer from genetically engineered Thermosynechococcus elongatus BP1 to wild-type E. coli DH5α. Gene 704: 49–58.

Warwick SI, Légère A, Simard MJ, James T. 2008. Do escaped transgenes persist in nature? The case of an herbicide resistance transgene in a weedy Brassica rapa population. Mol Ecol 17: 1387–1395.

Weaver M, Vedenyapina E, Kenerley CM. 2005. Fitness, persistence, and responsiveness of a genetically engineered strain of Trichoderma virens in soil mesocosms. Appl Soil Ecol 29: 125–134.

Yus E, Maier T, Michalodimitrakis K, van Noort V, Yamada T, Chen WH, Wodke Judith AH, Güell M, Martínez S, Bourgeois R, et al. 2009. Impact of Genome Reduction on Bacterial Metabolism and Its Regulation. Science 326, 1263–1268.

Zeng F, Qian J, Luo W, Zhan Y, Xin Y, Yang C. 2009. Stability of transgenes in long-term micropropagation of plants of transgenic birch (Betula platyphylla). Biotechnology Letters 32, 151.

Zhou J, Zhang H, Meng H, Zhu Y, Bao G, Zhang Y, Li Y, Ma Y. 2014. Discovery of a super-strong promoter enables efficient production of heterologous proteins in cyanobacteria. Sci Rep 4: 4500.

